# A Framework for Compressive On-chip Action Potential Recording

**DOI:** 10.1101/2025.09.26.678698

**Authors:** Pumiao Yan, Dante G. Muratore, E.J. Chichilnisky, Boris Murmann, Tsachy Weissman

## Abstract

Scaling neural recording systems to thousands of channels creates extreme bandwidth demands, posing a challenge for resource-constrained, implantable devices. This work introduces an adaptive, multi-stage compression framework for high-bandwidth neural interfaces. The system combines a Wired-OR analog-to-digital compressive readout with a digital core that adaptively requantizes, selectively samples, and encodes the neural signals. Although prior work suggests that action potential recordings can be re-quantized to approximately the signal-to-noise (SNR) number of bits without significantly degrading decoding performance, our results show that the required resolution can often be reduced even further. By matching the number of quantization levels to the electrode’s maximum SNR (⌈log_2_ SNR⌉ number of bits), we retain waveform fidelity while eliminating unnecessary precision that primarily captures noise. Recorded spike samples are selected using a mutual information-based criterion to preserve both spatial and temporal discriminative waveform features. A static entropy coder completes the pipeline with low computation overhead compression optimized for neural signal statistics. Evaluated on 512-channel macaque retina *ex vivo* data, the system preserves 90% of spikes while achieving a 1098× total compression over baseline.

## I. Introduction

**H**IGH-DENSITY neural interfaces capable of recording from thousands of neurons at single-cell resolution are transforming both neuroscience and clinical neurotechnology [1]. These interfaces provide access to fine-grained activity across large neural populations, which offer unprecedented insight into complex interactions between neurons and their cooperative behaviors [2]–[7]. Applications of neural interfaces range from basic studies of sensory processing to brainmachine interfaces for motor rehabilitation, vision restoration, and sensory augmentation [8], [9]. Meeting the demands of long-term, high-resolution *in vivo* recording requires systems that have many simultaneous recording channels and high temporal resolution, and must operate fully wirelessly for stable chronic use.

To support these applications, wired high-density microelectrode arrays (MEAs) with thousands of electrodes have been developed in research, enabling large-scale neural recordings with fine spatial and temporal resolution [10], [11]. However, existing fully implantable systems remain limited to roughly a thousand simultaneous channels [12] and are unable to preserve spike waveform information. As channel counts scale up, power consumption, silicon area, and data transmission requirements grow proportionally. Processing and transmitting this volume of data within the power and area constraints of a wireless implant becomes increasingly difficult. This motivates the need for hardware-friendly and power-efficient compression that reduces data through the data acquisition pipeline, without discarding critical spike-level information needed for downstream analysis.

In neural recordings, most of the information relevant to downstream processing is contained in the shape and timing of action potentials [13], [14]. These features enable essential tasks such as spike detection, spike sorting, and cell-type classification (the spike processing pipeline of such tasks is shown in Fig. 1). Spike detection identifies the timing and location of events, sorting clusters them by neuronal source, and cell-type classification associates units with specific cell types based on waveform and firing statistics. While the information required for each task varies (e.g., shape, amplitude, or timing), waveform shape remains a key feature, especially for more advanced analyses such as cell-type classification [15]– [17]. As a result, the structure of the spike waveform directly influences which features are retained for downstream interpretation and which can be disregarded as noise or redundancy.

**Fig. 1:**
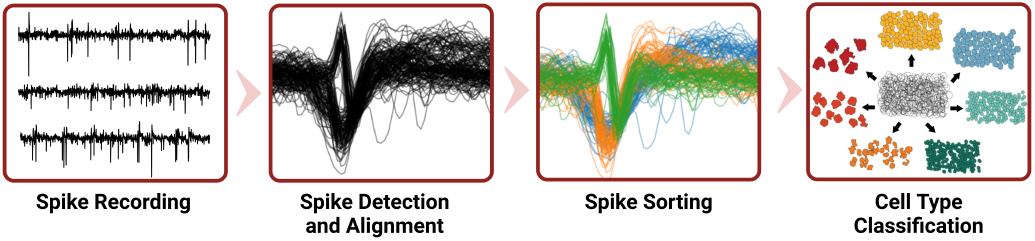
Action potential signal processing pipeline.

Conventional compression techniques typically rely on thresholding or temporal sub-sampling to reduce data volume [18]–[23]. While thresholding can be effective for simple decoding tasks, it discards waveform details needed for sorting and classification. Setting optimal thresholds also requires channel-specific tuning, which increases system complexity and power usage. Other approaches propose on-chip spike sorting [24]–[26] or compression following digitization [27]– [29], but digitizing full-resolution signals from thousands of channels remains a major bottleneck.

To avoid overwhelming bandwidth at any stage, compression must begin as close as possible to the analog front-end [30]. Ideally, this compression would be adaptive—preserving the relevant spike information for a given task while discarding non-informative baseline activity. As shown in Fig. 2, this requires rethinking the signal processing pipeline to integrate lossy compression at the analog-to-digital (A/D) interface.

**Fig. 2:**
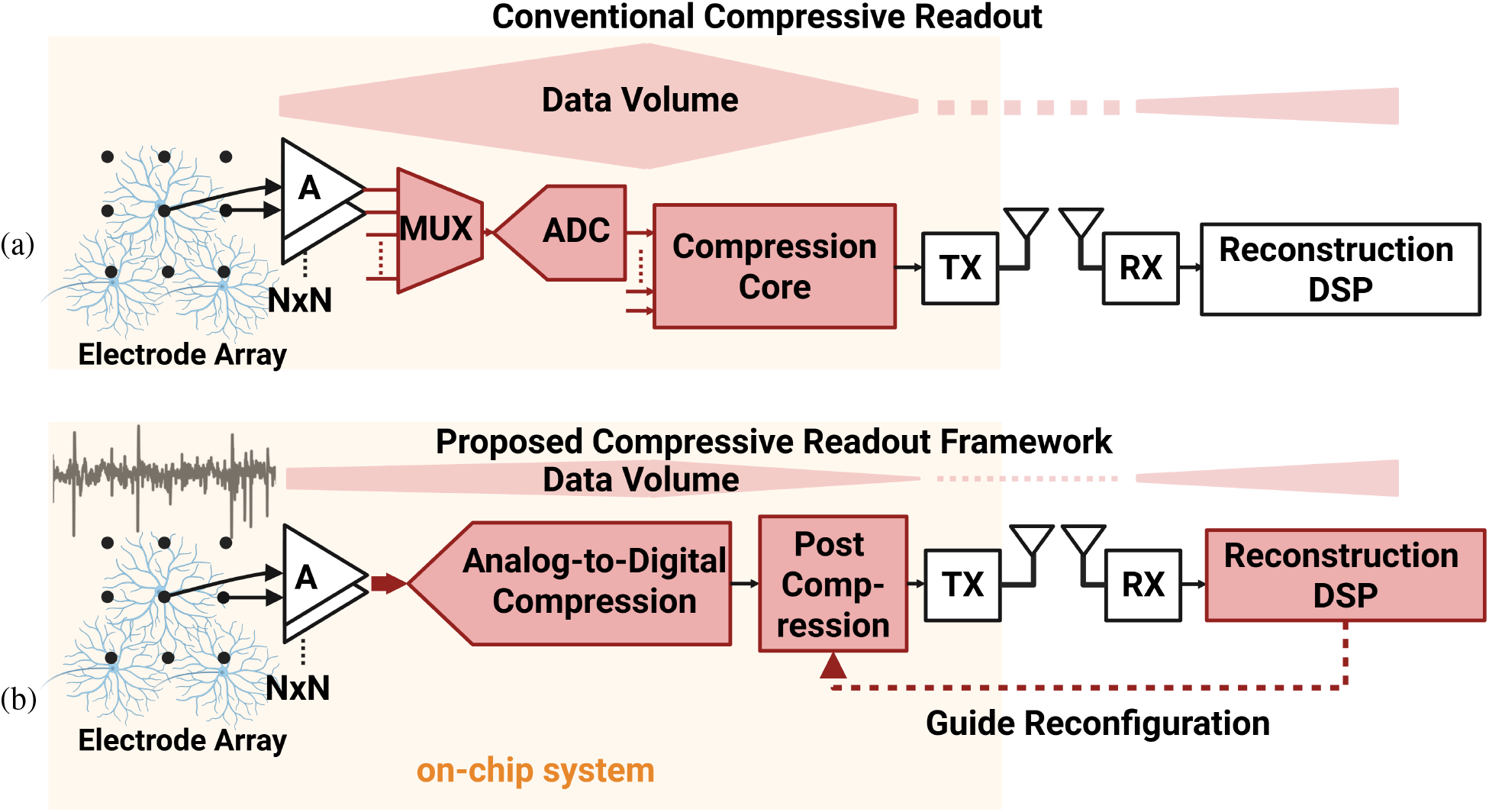
(a) Conventional and (b) proposed on-chip compressive readout framework.

In the conventional system architecture (Fig. 2(a)), signals from a multiplexed electrode array are first digitized in full by an ADC before undergoing any compression. This design imposes a heavy bandwidth and power burden at the front-end and requires ADCs that are capable of handling continuous high-rate data, including non-informative baseline samples. Compression occurs only after digitization, requiring more processing power and bandwidth.

Recent efforts have explored embedding spike sorting in hardware to enable low-latency, on-chip processing. For example, our prior work [31] presented a 1024-channel spike-sorting chip using event-driven detection and self-organizing map clustering, while Chen et al. [32] proposed an unsupervised geometry-aware sorting architecture. These approaches show promise, but typically focus on either spatial or temporal features, not both, and remain limited by power constraints.

In this work, we present a compressive readout framework that builds on previous analog-to-digital compression architectures [33], [34] and incorporates adaptive digital processing tailored to neural signals. As shown in Fig. 2(b), the system features a multi-stage pipeline that combines a Wired-OR analog-to-digital readout with an on-chip digital core that adaptively re-quantizes, selectively samples, and encodes spike waveforms. These operations are configured by an external module using signal statistics and waveform templates. Using this calibration-driven approach, the hardware applies compression strategies optimized for both data rate and signal fidelity.

We evaluate this framework through emulation on 512-channel *ex vivo* primate retina recordings. Results show that our system can preserve 90% of spike events – an improvement of 8 – 10% over state-of-the-art methods [31] – while achieving a 1098× total compression ratio over baseline recordings. These findings demonstrate that signal-aware, multi-stage compression can achieve substantial data reduction without sacrificing downstream performance.

This paper extends our prior work on Wired-OR compression by introducing and analyzing the full adaptive pipeline. Specifically:

- We develop a re-quantization strategy based on electrode SNR, eliminating excess quantization levels.
- We propose a mutual information–based selective sampling method that preserves both spatial and temporal waveform features.
- We implement a lightweight entropy coding scheme for efficient final-stage compression.
- We evaluate the pipeline on real 512-channel recordings and analyze performance across spike detection and sorting tasks.

The rest of this paper is organized as follows. Section II introduces the multi-stage compression framework, including the Wired-OR analog-to-digital readout and the adaptive digital post compression core. Section III presents simulation and experimental results using 512-channel neural recordings. Section IV discusses hardware implications and comparisons with existing methods. Section V concludes the paper.

## II. Multi-stage Compression Framework

We now present the implementation details of the proposed compressive readout framework, focusing on how compression is distributed across analog and digital domains to minimize bandwidth and power. The proposed system operates in two main phases: calibration and compression. During the Calibration Phase (Fig. 3a), a representative subset of recorded neural signals is analyzed offline to extract key statistical and structural features. Characteristic spike waveforms are identified, and relevant metrics such as SNR, firing rates, and salient waveform features are estimated. These metrics guide the selection of optimal compression parameters, which are loaded into the on-chip digital compression core. Tailoring the system parameters to the underlying neural activity in this way ensures that compression decisions, such as quantization levels or sampling intervals, are application-aware and reflective of the unique signal properties observed during calibration.

**Fig. 3:**
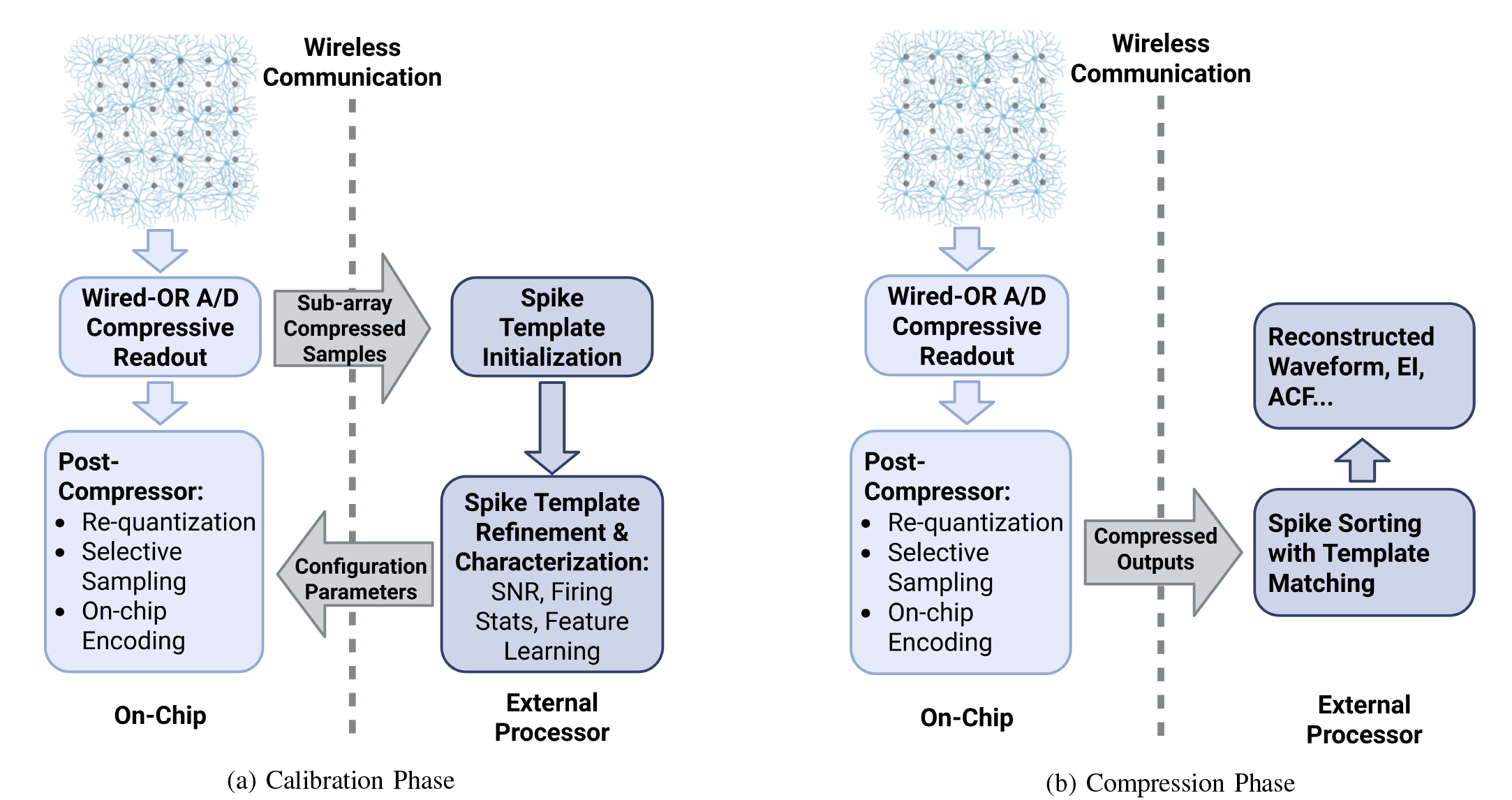
On-chip compressive readout framework.

Once the calibration parameters have been established, the system transitions into the Compression Phase (Fig. 3b). Neural signals are continuously acquired, and the first level of compression occurs immediately at the analog front-end through the Wired-OR A/D compressive readout. This mixed-signal architecture exploits the sparsity of neural activity: spike-related voltages, which deviate sharply from baseline and are less frequent, are more likely to be uniquely encoded, while baseline-level voltages, being more frequent, are prone to collision in the row and column readout and naturally suppressed. As a result, the data volume is substantially reduced while preserving nearly all spike-relevant samples, alleviating pressure on downstream processing. Following this mixed-signal compression, a digital post-compression core refines the data further through re-quantization, selective sampling of spike waveforms, and low-overhead on-chip encoding. This multi-stage strategy ensures that each subsequent compression step operates on a smaller data stream, thereby reducing both power consumption and bandwidth requirements without sacrificing essential spike information.

### A. Wired-OR analog-to-digital compressive readout

The Wired-OR analog-to-digital (A/D) compressive readout [33]–[35] (as shown in Fig. 2(b)) serves as the first stage of the proposed multi-stage compression framework, directly reducing data at the A/D interface. This hardware-efficient and event-driven approach acquires neural data while simultaneously performing data compression and channel multiplexing. Each pixel samples the input voltage and encodes it into a pulse position based on a globally distributed ramp. Pulses from multiple pixels are merged on shared wires using Wired-OR circuitry. If only one pixel fires at a given time step, both its location and voltage level can be uniquely decoded (as shown in Fig. 4(b)). If multiple pulses coincide—i.e., a collision occurs—the data is discarded, achieving compression by exploiting signal sparsity (see Fig. 4(c-d). This technique exploits the sparsity and diversity of neural signals to minimize redundant sampling and digitization while maintaining the key features required for downstream analysis.

**Fig. 4:**
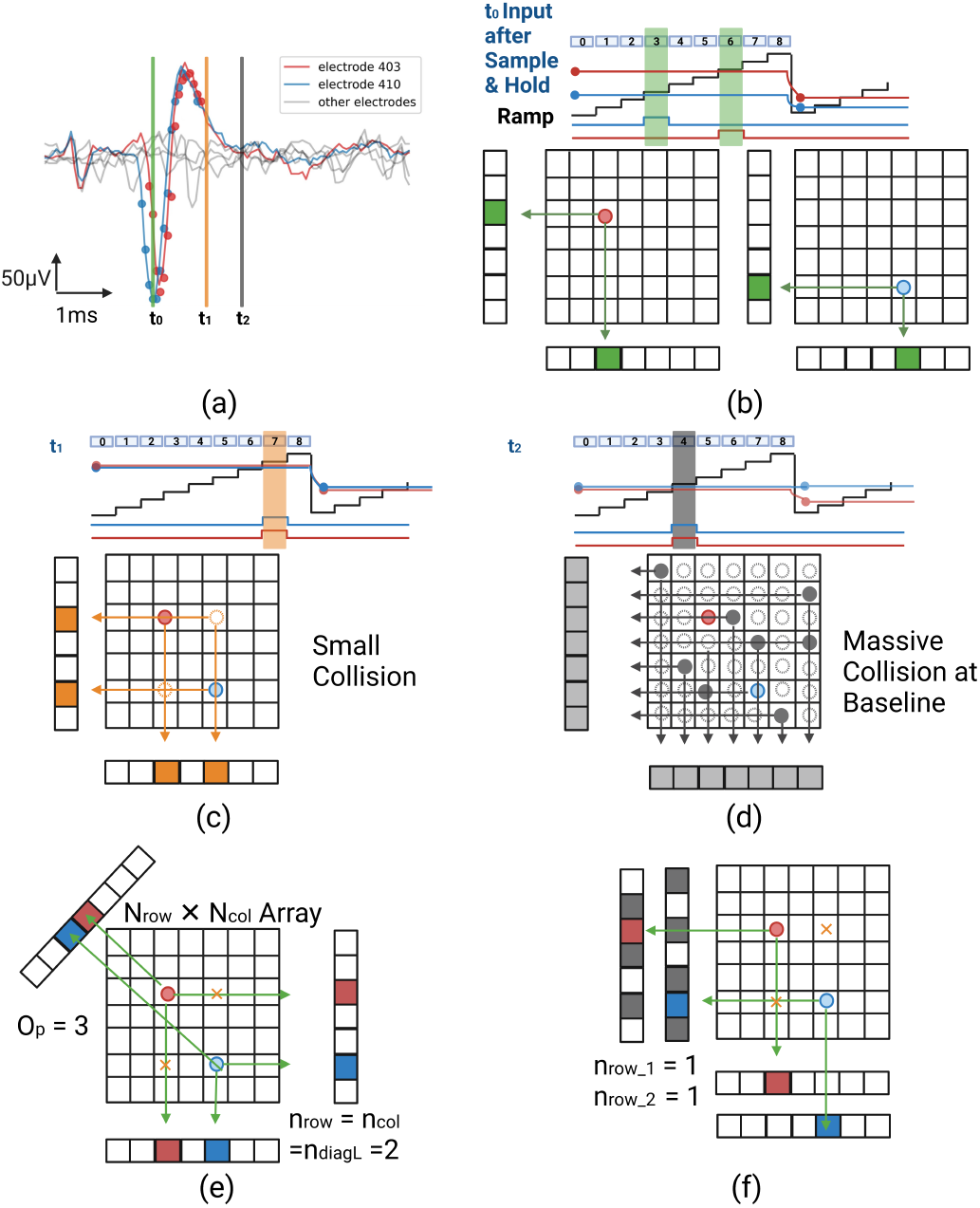
Wired-OR readout concept [34]. (a) A snippet of action potential waveform seen on different electrodes. (b) Conversion of voltage to pulse position and collision-free readout of one pixel. (c) A collision between two pixels. (d) Massive collision across the array at the baseline level. (e) Diagonal wiring conceptual drawing. (f) Interleaving wiring.

Unlike conventional A/D architectures that sample continuously across all channels, the Wired-OR readout refrains mostly spike-related events, significantly reducing data rates and computational overhead. This spike-driven encoding process ensures that only the most relevant neural activity is retained, making the system highly scalable and power-efficient for brain-machine interface applications. Due to their temporal and spatial sparsity, spike-related samples are more likely to be unique, they are retained, while baseline-level voltages—which are more frequent and thus prone to collision—are naturally filtered out. Wiring strategies such as diagonal [34] and interleaved [33] layouts can help resolve small collisions and recover more spike information, also shown in Fig. 4(e-f).

By pre-compressing neural data at the analog interface, the Wired-OR readout reduces the data bandwidth burden on subsequent processing stages by 15 ∼ 50 ×, depending on the wiring strategy and neural firing rate. This efficiency was validated in ex vivo experiments using a taped-out chip, where compression rates ranged from 111.2× (single-wire) to 38.8× (four-wire) configurations [35]. The digital compression core, detailed in Section II-B, further refines the signal through re-quantization, selective sampling, and on-chip encoding, ensuring optimal balance between compression efficiency and neural signal fidelity.

### B. Digital compression core

The output of the Wired-OR readout stage is a sparse set of events, each represented by its row and column address along with the amplitude (encoded in ADC counts). These elements - spatial address and amplitude - form the input to the post compression core, which aims to further reduce data size by keeping critical information available for downstream analyses. The digital core first treats address and amplitude data as separate streams, optimizing compression strategies tailored to each. For the amplitude, we explore re-quantization to a lower resolution, leveraging the insight that many spikes can be accurately reconstructed at a coarser bit depth. In addition, selective sampling is applied to discard redundant or low-information spike samples. The remaining critical samples are then passed through a lightweight entropy coding stage inspired by Huffman coding. Instead of performing full encoding in hardware, we use a precomputed lookup table that approximates optimal codes based on amplitude and spatial statistics gathered during the calibration phase. This hybrid scheme achieves high compression ratios while maintaining hardware simplicity and preserving spike timing and shape characteristics essential for downstream decoding and sorting.

#### 1) Re-quantization

State-of-the-art neural interface systems are equipped with ADCs of 10-16 bit resolution [10], [12], [36]–[39] driven by multiple considerations such as signal drift, motion artifacts, and dynamic range. Although prior work [19] suggests that action potential data can be requantized to about the SNR number of bits without significantly degrading decoding performance, our results indicate that the required re-quantization resolution can often be reduced even further. In an electrode of a MEA, the measured voltage *V* (*z, t*) can be modeled as [40]:

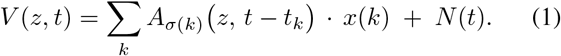

Here, *z* denotes the electrode location, *t* is the continuous time, and *k* indexes distinct neurons or clusters. Each cluster *σ*(*k*) has an associated waveform template *A*_*σ*(*k*)_, shifted by the spike time *t*_*k*_ and scaled by the spike amplitude *x*(*k*). The term *N* (*t*) captures additive noise present in the recording.

For a spike event, the SNR is approximated by [36]:

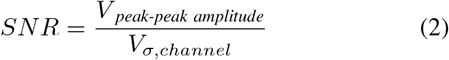

The amplitude of the spike peak-to-peak (*V*_*peak-peak amplitude*_) is determined by identifying the electrode with the most significant peak-to-peak difference for each spike. The noise level (*V*_*σ,channel*_) is calculated as the median absolute deviation when no action potential is detected on the electrode.

Intuitively speaking, if we look at the recording of spikes on a certain electrode, the observed voltage level is the summation of the underlying noise-free signal and noise that can be modeled as a Gaussian distribution (see Fig. 5). When the electrode records spikes only up to a certain *SNR*_*max,electrode*_ (following the definition of Eq. 2), the quantizer only needs to differentiate *N*_*steps*_ quantization levels, where:

**Fig. 5:**
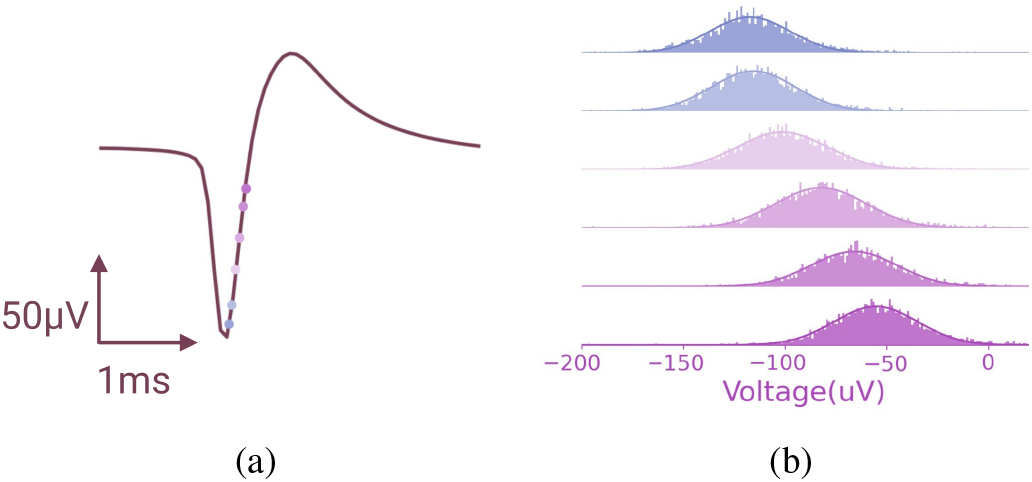
Empirical demonstration of additive Gaussian noise in extracellular action potential recording. (a) Average spike waveform of a given cell-electrode pair. (b) Real, measured voltage distribution at a specific sample point across multiple spike occurrences.

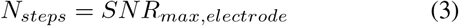

because finer resolution would merely differentiate noise. Hence, the minimum bit depth required is:

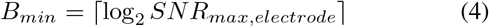

From a theoretical perspective, this interplay between universal filtering and lossy compression aligns with established results in information theory and signal processing. The foundational works of [41]–[43] have characterized noise filtering via data compression and indirect rate-distortion problems, respectively, underscoring the effectiveness of compression-based denoising. Additionally, the principles articulated in [44], [45] reveal that compressing noisy data can inherently facilitate noise reduction when the compression is optimized for a distortion tuned to the level and characteristics of the noise. Practical manifestations of this approach are extensively discussed in prior research [44], [46], reinforcing the viability of compression-driven filtering methods in real-world scenarios.

Further justification for our use of uniform scalar quantization followed by universal filtering arises from both theory and practice. It is well-established that, under squared-error distortion and in the presence of additive Gaussian noise, uniform scalar quantization achieves near-optimal performance [47]. This setting applies directly to neural signals, which are commonly modeled as clean spike waveforms corrupted by zero-mean Gaussian noise (see Fig. 6). Prior work [47] has shown that quantizing the noisy signal using a quantizer designed for the clean source yields distortion nearly matching that of the optimal scheme. Thus, uniform scalar quantization not only simplifies implementation by obviating the need for optimized quantizer thresholds but also incurs minimal performance loss^1^.

**Fig. 6:**
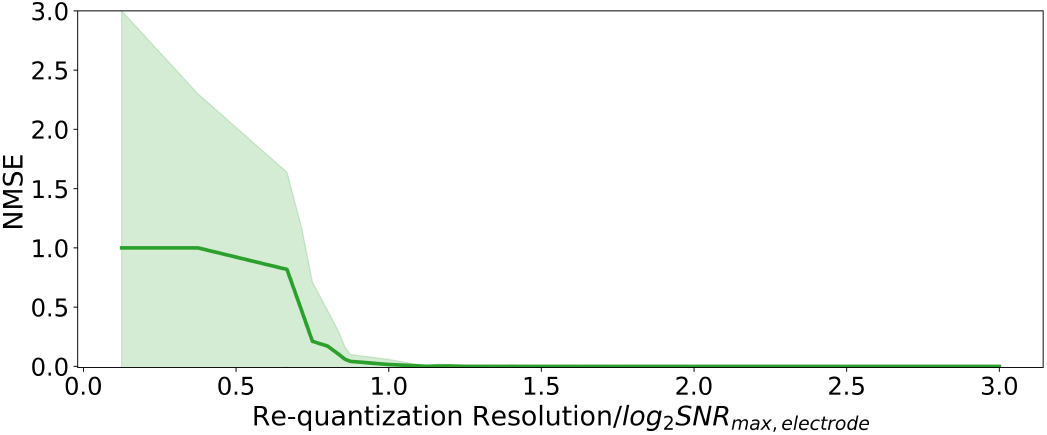
Normalized mean square error (NMSE) versus requantization resolution, where the x-axis is expressed as the ratio of each electrode’s chosen bit depth to *B*_min_.

An experiment of requantizing the action potential recording from an originally 12-bit quantized dataset is conducted. The requantization resolution for each electrode in three 512-channel primate *ex vivo* retina recording datasets is normalized to the proposed *B*_*min*_ defined by Eq. 4. As shown in Fig. 6, the green curve shows how NMSE^2^ decreases sharply as the bit depth approaches or exceeds *B*_*min*_, indicating that the requantization error can be substantially mitigated by allocating just enough bits to match the electrode’s maximum SNR-based bits. The shaded region depicts the variability across different channels, illustrating the NMSE is consistently small across electrodes with a requantization resolution of *B*_*min*_.

We also studied the average spike waveform distortion introduced by re-quantizing with *B*_*min*_ bits resolution across different cell-electrode pairs where spikes have a wide range of SNR. Fig. 7 shows that for *SNR* ≥ 5, less than 1% of NMSE is introduced by re-quantization.

**Fig. 7:**
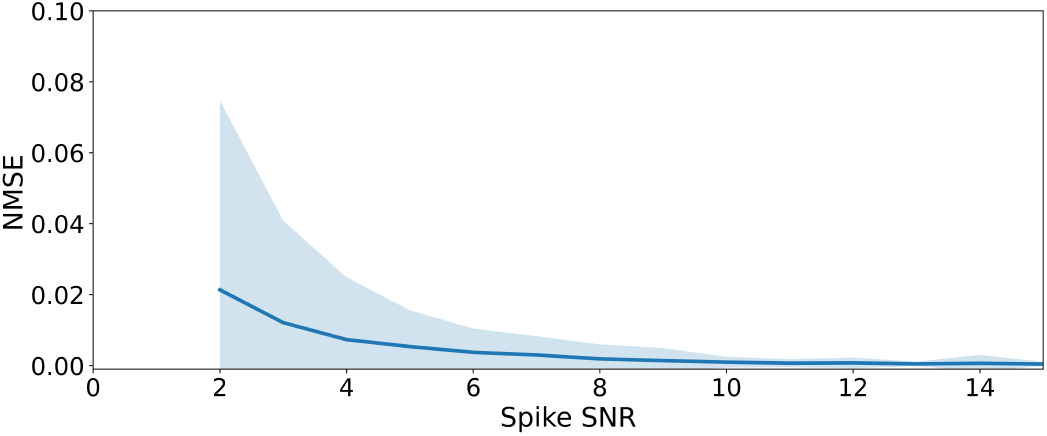
Normalized Mean Square Error (NMSE) versus spike SNR for re-quantization at *B*_*min*_ bits.

During the calibration phase (Fig. 3a), a single sweep across all channels is performed to extract signal statistics used to estimate the SNR and determine *B*_*min*_ for each channel. In this mode, full waveforms from each electrode are sampled and digitized without thresholding or compression, enabling accurate baseline noise characterization.

Because SNR can vary over time due to factors such as electrode impedance changes, tissue response, or other nonstationary effects, the system includes a recalibration mechanism. It periodically re-enters calibration mode to update noise estimates and recompute *B*_*min*_. The re-calibration interval is empirically determined based on the application, as different recording modalities (e.g., retinal vs. intracortical) exhibit different SNR dynamics. This adaptive calibration ensures that the quantizer remains well-tuned to current signal conditions, preserving both compression efficiency and signal fidelity.

#### 2) Selective sampling

Following the compressive Wired-OR readout and digital re-quantization, further compression gains can be realized by reducing the number of time samples retained for each spiking event. Rather than transmitting all of the Wired-OR outputs, we identify a small set of time points that carry the most discriminative information across neural units. This selective sampling process eliminates redundant or less-informative samples, preserving only those samples essential for downstream tasks such as spike sorting or neural decoding. Selective sampling is configured during the Calibration Phase, where informative samples are identified off-chip based on statistical differences across spike waveforms. The selected sample indices are then programmed into the on-chip digital core to enable compact spike representation during realtime data streaming.

Our approach is motivated by information-theoretic principles, yet implemented using lightweight operations suited to streaming. The procedure unfolds in three stages: an initial unsupervised clustering, mutual information-based sample ranking, and optional online refinement.

##### a) Unsupervised Clustering Initialization

We first start with performing an initial unsupervised clustering on the full Wired-OR spike waveforms. Here we use k-means clustering (with *k* set to a conservative number of 5, empirically chosen). This provides provisional cluster assignments without any feature selection bias.

Given spike waveforms of length 61^3^ (each approximated as a true spike shape plus Gaussian noise), we need to find which time sample indices are most informative for distinguishing unknown clusters (typically 3–5 clusters). In an unsupervised scenario (no prior labels), we seek statistically sound, efficient feature-ranking methods that highlight time points contributing most to cluster separation. We focus on global importance across all clusters seen on the electrode (not per-cluster specific) and avoid heavy machine-learning models. Below, we outline the principle of our method. We also discuss their computational efficiency and how they can fit into a real-time feature selection pipeline.

##### b) Information-Theoretic Sample Selection

While mutual information between each sample requantized value *X*_*t*_ and the unit identity *C* is an ideal theoretical measure of importance,

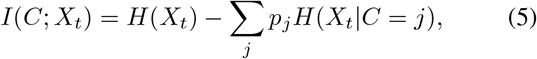

its computation is costly and ill-suited to real-time or low-power implementations. We therefore approximate this metric under the Gaussian noise model (see Eq. 1), where mutual information simplifies to a ratio of variances:

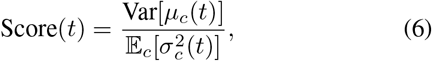

with 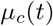 as the mean waveform value at time *t* for cluster *c*, and 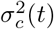 as the within-cluster variance. This discriminability score captures the same intuition: samples with high inter-unit variability and low intra-unit noise are more informative for separating spikes from different sources.

We compute this score for each time index and retain only the top 3–5 samples with the highest discriminability, forming a compact, information-rich representation of each spike. While we do not claim this number to be theoretically optimal, it was found to be empirically sufficient across multiple datasets. This choice reflects a practical tradeoff between reducing input dimensionality and preserving clustering performance.

##### c) Refined Clustering and Online Update

After feature selection, clustering is rerun using the reduced feature vectors. This not only accelerates computation but can improve accuracy by suppressing noisy and less-informative samples. To support long-term use in dynamic recording environments, the discriminability scores can be updated incrementally as a subset of new spikes, with all Wired-OR spike samples, are periodically transmitted. We maintain running estimates of per-cluster means and variances at each time point, enabling online updates to feature rankings with minimal overhead.

These statistics converge quickly and require only *O*(*k*) computation per spike, allowing the system to adapt to changing neural conditions while maintaining compression efficiency.

#### 3. On-chip encoding

To efficiently encode remaining samples, we implement a Huffman-inspired entropy coder using a precomputed lookup table. During the system’s calibration phase, histograms of address and amplitude values are collected and used to build codebooks optimized for the neural signal distribution. The codebook is refreshed at each recalibration, keeping the coding efficiency near the true entropy of the evolving signal distribution.

Instead of constructing Huffman trees at runtime, which is expensive in both time and silicon area, we generate static Huffman tables offline and store them on-chip. These tables translate each sample’s spatial and amplitude value into a fixed-length or variable-length code via a simple lookup. This method eliminates the need for real-time histogramming or adaptive coding logic, enabling ultra-low-power, high-speed compression.

The channel coordinates and spike amplitude exhibit skewed distributions due to structured neural activity, allowing Huffman coding to achieve substantial gains. Coupled with selective sampling and re-quantization, this final stage completes the multi-step pipeline from raw events to fully compressed and hardware-ready output.

## III. Simulation and Results

To evaluate the performance of our multi-stage compression framework, we conduct experiments on 512-channel *ex vivo* primate retina datasets recorded at 20kHz. These datasets are processed in software to emulate the full pipeline, including Wired-OR analog-to-digital compressive readout, requantization, selective sampling, and entropy coding. Each stage is designed to reduce the total data volume while preserving critical information for downstream spike processing. Parameter recalibration was performed every 10 minutes, which was sufficient to maintain stable compression performance over time. We further benchmark our framework against state-of-the-art on-chip spike sorters [25], [31] using a publicly available multi-channel Neuropixels dataset [48].

Fig. 8 provides a detailed example of the full pipeline applied to one channel and its neighboring electrodes. The top row (see Fig. 8 (a–c)) shows the extracted electrical images (EIs^4^) of units recorded on electrode 501 in the array, initial spike templates obtained through unsupervised k-means clustering, and the panel (c) shows the refined templates obtained after re-clustering using only the discriminative samples selected during the selective sampling phase, as described in the refined clustering and online update stage. This step enhances both accuracy and efficiency by removing noisy or redundant dimensions from the feature space. The bottom row highlights the discriminability scores computed for two adjacent electrodes: Fig. 8 (d) for electrode 501 and Fig. 8 (e) for electrode 502. Notably, in this example, electrode 502—despite not being the primary site of the spike—exhibits more peaked and informative samples for class separation. This underscores the value of spatially informed selective sampling, where nearby electrodes can provide higher-discriminability features than the electrode with the largest spike amplitude.

**Fig. 8:**
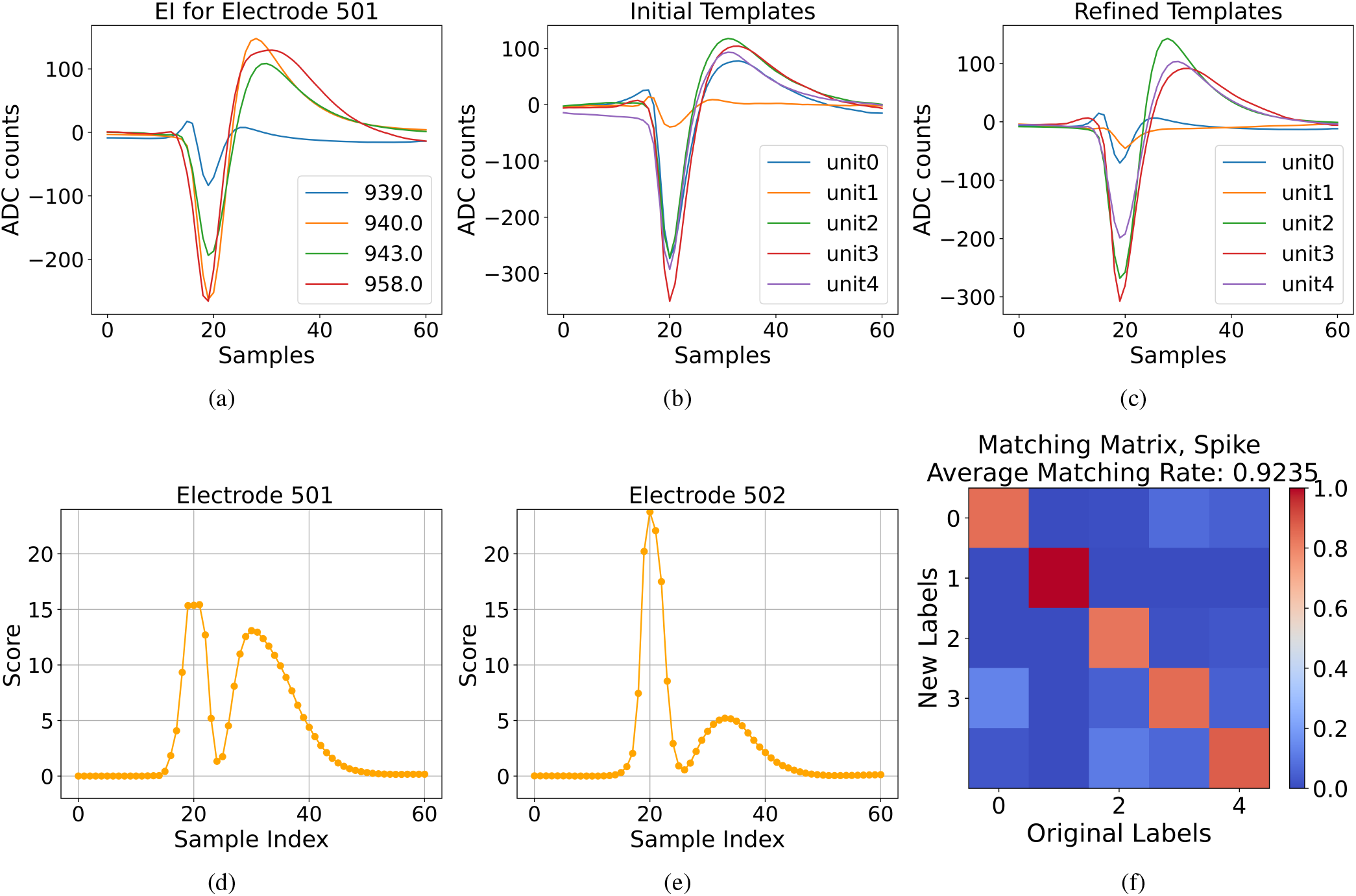
An example of processing through electrode 501 in simulation. (a) Ground truth electrical image (EI) of the cells on electrode 501. (b) Initialized templates by clustering the detected and aligned events. (c) Refined and updated templates after iterating through the dataset. (d) Discriminability score of electrode 501 given the collected statistics after the calibration phase. (e) Discriminability score of electrode 502, which is near 501. (f) The matching matrix of the final clustering results and ground truth labels (different cells with the largest spike on electrode 501 and “garbage”).

Fig. 8 (f) shows a spike matching matrix between the clustering output from compressed data and ground truth unit labels. Clusters are aligned based on their dominant electrode. To address duplicate clusters across nearby electrodes, we apply a merging procedure based on the Hierarchical Adaptive Means (HAM) clustering strategy [49], which compares intercluster distances against a noise-informed threshold and consolidates redundant units in a hardware-efficient manner. The resulting average matching rate in this example is 92.35%.

To assess generalization beyond a single electrode, we evaluated the percentage of spikes preserved across the entire dataset. Fig. 9 summarizes spike recall as a function of signal-to-noise ratio (SNR) for different cell-electrode pair in the dataset. We compare two representative schemes: (1) only compress with Wired-OR a 1-wire 6-bit ramp configuration, achieving a compression ratio (CR) of 274.35, and (2) the full compression pipeline with a 10-bit ramp, eight interleaved wiring Wired-OR configuration, and the addition of digital post-compression, resulting in a total CR of 1098.23.

**Fig. 9:**
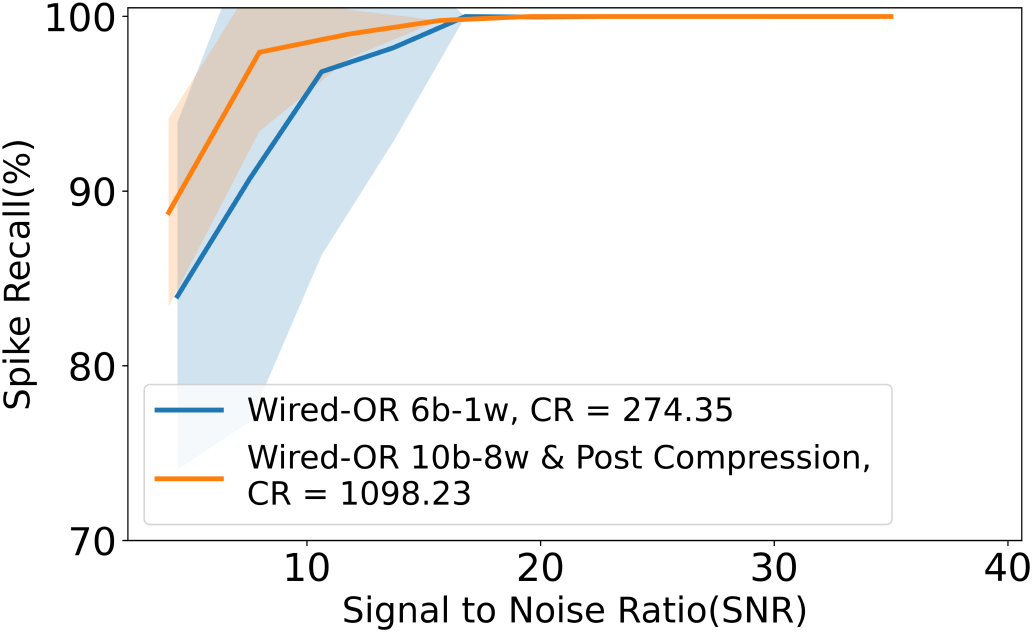
Spike recall for ExVivo-1, across varying SNRs. The full pipeline achieves higher recall at higher compression.

As shown in Fig. 9, both configurations achieve high spike recall across a broad SNR range, with the full compression pipeline consistently outperforming the baseline Wired-OR-only setup. Notably, even under aggressive compression, spike recall exceeds 95% for moderate SNR values (SNR ≥ 8) and approaches 100% for higher-quality signals. The shaded region represents variability across electrodes for different SNR ranges, reflecting electrode-specific SNRs.

These results demonstrate that combining Wired-OR read-out with post-compression stages—including re-quantization, selective sampling, and Huffman-inspired encoding—can significantly increase compression efficiency without compromising spike detection fidelity.

We next evaluate how our proposed compression framework compares against other spike compression and sorting approaches [31] on the same 512-channel *ex vivo* primate retina datasets (referred to hereafter as ExVivo-1 and ExVivo-2) and Neuropixel dataset [48]. These datasets are chosen as representative cases of neural recordings with varying SNR distributions, which significantly affect compression performance—particularly in the re-quantization and entropy coding stages. Specifically, ExVivo-1 has a mean SNR of 16.16 with a standard deviation of 6.99, while ExVivo-2 exhibits a mean SNR of 12.64 with a narrower spread (standard deviation of 6.06).

Fig. 10 shows the spike clustering accuracy across three datasets using our method (with information-theoretic selective sampling), the spatial spike sorting method reported from [31], and a temporal spike sorting baseline.

**Fig. 10:**
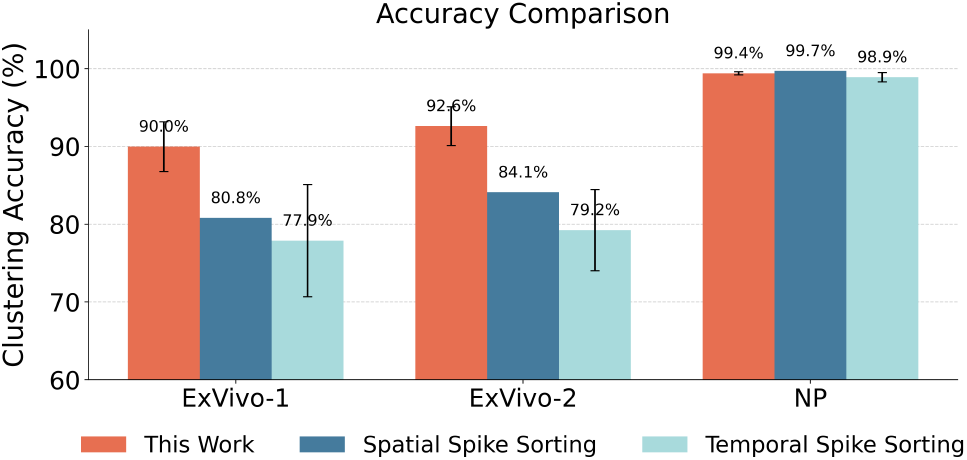
Spike clustering accuracy across two retina datasets and one public Neuropixel dataset comparing three methods: our work using discriminability-driven selective sampling, the spatial spike sorting method proposed in [31], and a temporal single-electrode baseline. Our method achieves the highest average accuracy and more consistent performance across units.

The temporal method performs spike sorting independently on each electrode by using all Wired-OR spike samples and linearly interpolating missing ones before classification, following the methods that we previously proposed in [50]. The spatial method, as proposed in [31], combines spikes across electrodes and leverages spatial correlation for clustering using a self-organizing map approach. While the spatial method shows strong performance, it does not report the variability of sorting accuracy across units. Our method not only achieves the highest mean clustering accuracy in both datasets (90.0% for ExVivo-1, 92.6% for ExVivo-2 and 99.4% for Neuropixel data), but also shows lower variability compared to using only temporal features, indicating robust performance across all neural units.

To understand how each stage of the compression pipeline contributes to overall compression, we analyze the normalized data size after each component. As shown in Fig. 11, the Wired-OR stage alone achieves 35× and 19× compression on ExVivo-1 and ExVivo-2, respectively. With selective sampling and re-quantization, an additional 11× reduction is observed, followed by a further 3× from Huffman-inspired entropy coding. The overall compression rate exceeds 1000×, with acceptable loss in spike fidelity. In the Neuropixels dataset [48], the lower SNR (∼ 7) and long-shank electrode architecture result in significantly greater compression gains—achieving 358 ×from re-quantization and selective sampling, and an additional 4.8× from Huffman-inspired coding. Notably, the compressibility in the selective sampling stage depends on the number of retained samples; in this analysis, we select the top 5 most discriminative time points per spiking event on an electrode and its neighboring electrodes.

**Fig. 11:**
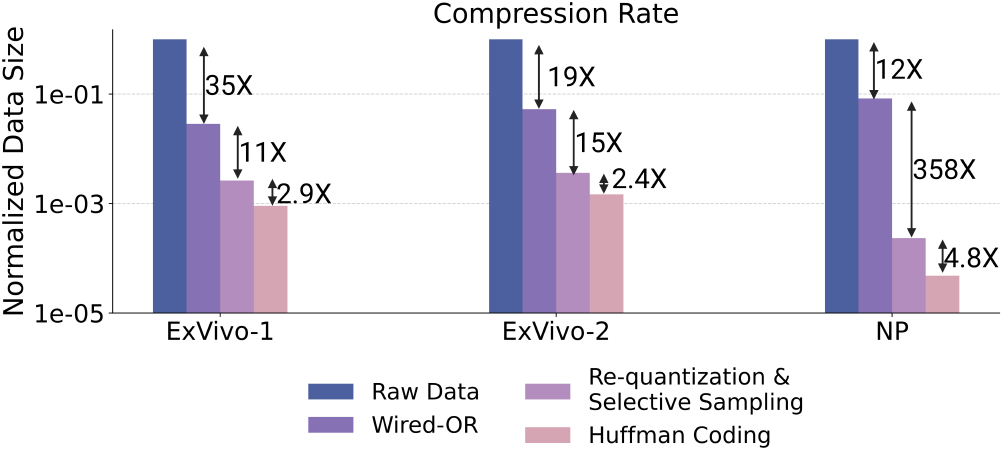
Compression rate comparison across pipeline stages for three datasets. Starting from raw data, the Wired-OR stage provides the first major reduction. Re-quantization and selective sampling yield another 11 ×, followed by Huffman coding with 2.5× further reduction. Each dataset retains 5 informative samples per spike.

## IV Discussion

### A. Implementation and benchmarking of digital compression core

Fig. 12 shows the proposed hardware architecture of the digital compression core that follows the Wired-OR analog-to-digital compressive readout. The system is divided into three main stages: (1) re-quantization, (2) discriminability-driven feature selection, and (3) approximate entropy coding via a lightweight Huffman encoder.

**Fig. 12:**
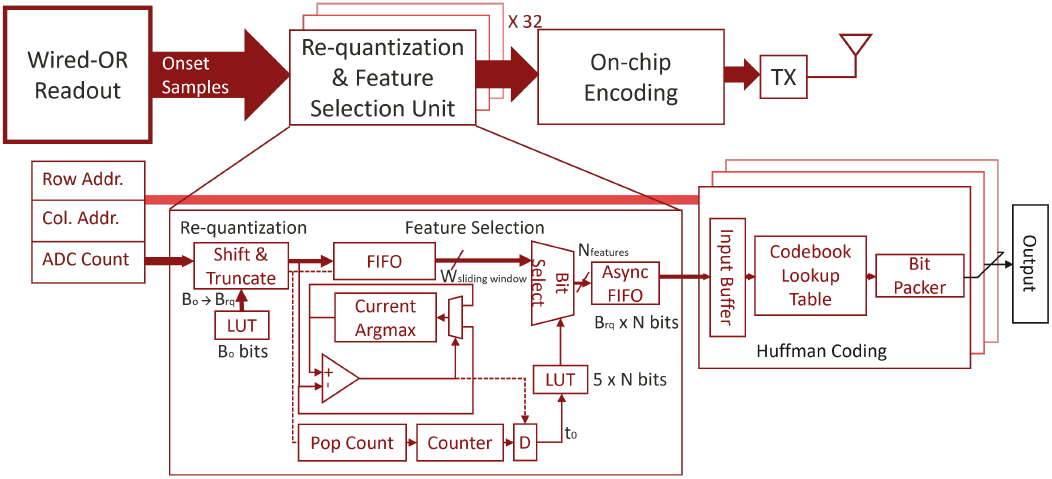
Hardware architecture of the proposed digital compression core. The system receives sparse spike samples from the Wired-OR stage and performs re-quantization, sample selection, and Huffman-style encoding.

Incoming spike events from the Wired-OR stage are represented as sparse triplets: row address, column address, and ADC count. These are first fed into the “Re-quantization and Feature Selection Unit”, which handles both amplitude compression and sample reduction.

In the re-quantization block, a programmable lookup table (LUT) determines the quantization resolution (*B*_*o*_ → *B*_*rq*_, where *B*_*o*_ is the length of the Wired-OR amplitude output, *B*_*r*_*q* is the reduced bit width after re-quantization) based on the maximum SNR per channel. A simple shift-and-truncate operation compresses the amplitude values. This enables amplitude-specific precision tuning across electrodes with negligible computation overhead.

For sample selection, incoming spike waveforms are buffered in a FIFO and processed using a sliding window. A lightweight argmax engine computes the spike peak timing, and the most informative *N* samples are selected. The selection is driven by a second LUT that is trained offline during a calibration phase. Selected samples are forwarded via an asynchronous FIFO to the encoder.

Finally, the data passes through an on-chip Huffman encoder that uses a fixed codebook precomputed during the calibration phase. The encoder maps frequent sample patterns to shorter binary representations using a lookup table, and a bit packer serializes the output. This design enables approximate entropy-coded compression with low hardware complexity and avoids the need for runtime codebook updates.

To assess implementation feasibility, we provide an estimated breakdown of complexity and energy consumption for the digital compression core. Table I summarizes gate count^5^ and energy cost per spike processed for each major module, based on standard logic primitives and published energy models for 28nm CMOS [51], [52]. Gate counts are estimated by decomposing each module into standard digital building blocks such as flip-flops, adders, comparators, and control logic, using representative gate-equivalent costs derived from synthesis reports and textbook implementations [53], [54].

**Table I.**
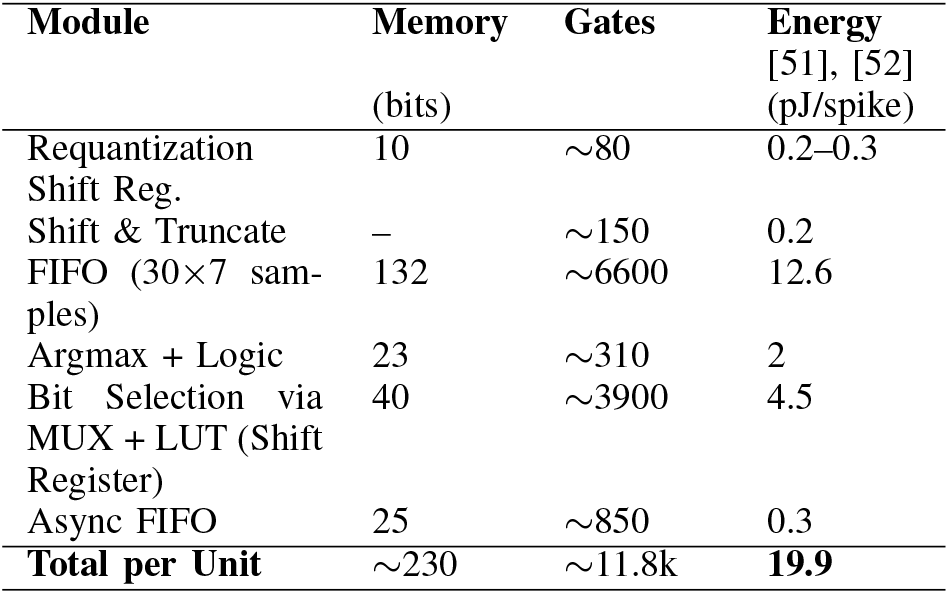
Estimated Memory, Gate Count, and Energy per Re-quantization and Feature Selection Unit (28nm LP)

The entire post-compression system is designed for shared processing across channels. A single Requantization & Feature Selection Unit is time-multiplexed across multiple channels. Based on spike event rates modeled empirically, only 32 such units are required to support a 1024-channel array. This architecture reduces the power and area overhead compared to per-channel buffering, while preserving the low-memory footprint that is a key advantage of the original Wired-OR design.

Including a conservative memory estimation of the Huffman lookup table of 4KB^6^, the total on-chip memory required for the post-compression stage across 1024 channels is approximately 4.92 KB. The entire logic complexity is well within feasible bounds for modern implantable SoCs.

#### a) Power Savings

Assuming a baseline wireless transmission energy of 72.9*pJ/bit* [55], the energy cost per bit after compression is broken down as follows^7^:

- 0.066*pJ/b* from re-quantization and selection,
- 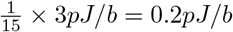 from Huffman encoding [56],
- 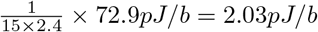 from transmission.

Total energy per transmitted bit: 2.296*pJ/* compared to transmitting all of the original Wired-OR outputs, this corresponds to an additional 31.75× power reduction.

This analysis confirms that the post-compression stage delivers significant energy and bandwidth benefits with low hardware overhead, making it highly suitable for implantable or portable neural interface systems.

### B. Comparison with Prior Work

Table II compares the proposed compression framework to previous neural signal compression and spike sorting approaches. Compared to conventional thresholding-based spike detection [21], which incurs low overhead but discards waveform detail, we maintain full waveform shape and achieve 3–10× higher compression compared to this reported thresholding approach. While ML-based autoencoders [27], [29] can offer similar compression ratios, they require training, inference, and high memory resources—making them less suitable for ultra-low-power edge systems. On-chip spike sorting techniques [25], [31] extract spike times and unit labels in hardware, often discarding waveform information, and typically rely on fixed temporal or spatial features. In contrast, our system supports tunable feature selection via an information-theoretic approach and retains waveform fidelity. Among all methods surveyed, Wired-OR is the only one to support high compression, waveform preservation, and scalable implementation without requiring large memory, floating-point operations, or machine learning. This makes it particularly well suited for large-scale neural recording systems targeting real-time closed-loop applications.

**Table II.** Comparison With Prior Work.

| Compression Approach | Thresholding Spike Detection [21] | ML-based Autoencoder [27] | Neuromorphic Compression [23] | On-chip Spike Sorting with Temporal Feature [25] | On-chip Spike Sorting with Spatial Feature [31] | Wired-OR + Post Compression (This Work) |
| --- | --- | --- | --- | --- | --- | --- |
| Memory Required | 1kB | 18kB | N/A | 98.08kB | 14kB | 4.92kB |
| Low Computation Overhead | ✓ | 79.25K MACs per spike | ✓ | x | ✓ | ✓ |
| No. of Channels | 64–256 | 256 | 384 | 96–384 | 128–1024 | 512 |
| Preserves Waveform | x | ✓ | ✓ | x | x | ✓ |
| Reported Accuracy | 97.4% | N/A | 92% | 84.3%–98.7% | 80.8%–99.7% | 90%–99.4% |
| Compression Ratio | 10–116× | 20–500× | 20–200× | 240–39272× | 1575–177768× | 678–20621× |

## V. Conclusion

We have presented an end-to-end framework for adaptive compressive neural signal acquisition and digital compression. Building on our prior work on Wired-OR analog-to-digital compressive readout [33], [34], this paper introduces a post-compression architecture that combines re-quantization, selective sampling based on spike discriminability, and hardware-friendly entropy encoding. Together, these components form a scalable system that dramatically reduces data rates while preserving spike information critical for more fine-tuned spike sorting algorithms.

Evaluated on 512-channel *ex vivo* primate retina recordings, the system achieves over 1000× compression while preserving 90% spike sorting accuracy. The design balances spatial and temporal feature retention with hardware efficiency, and supports runtime adaptation to varying signal conditions.

## VI. Biography Section

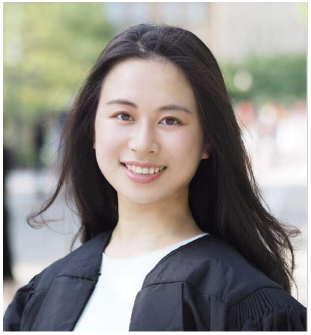

**Pumiao Yan** received the B.Sc. degree in Electrical and Computer Engineering from Cornell University in 2018, and the M.S. and Ph.D. degrees in Electrical Engineering from Stanford University in 2020 and 2025, respectively. She was a Seth A. Ritch Bio-X Graduate Student Fellow during her Ph.D. studies. Her research focuses on algorithm-hardware co-design and signal processing for analog-to-digital compression hardware architectures for neural interfaces.

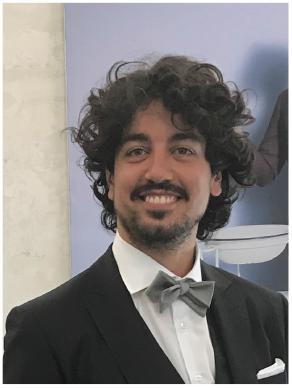

**Dante G. Muratore** is an assistant professor in the Microelectronics department at the Delft University of Technology. He received the B.Sc. degree and the M.Sc. degree in Electrical Engineering from Politecnico of Turin in 2012 and 2013, respectively. He received the Ph.D. degree in Microelectronics from the University of Pavia in 2017 in the Integrated Microsystems Lab. From 2015 to 2016, he was a Visiting Scholar at Microsystems Technology labs at the Massachusetts Institute of Technology. From 2016 to 2020, he was a Postdoctoral Fellow at Stanford University. He is the recipient of the Wu Tsai Neurosciences Institute Interdisciplinary Scholar Award. His research focuses on hardware and system solutions for high-bandwidth brain-machine interfaces that can interact with the nervous system at natural resolution.

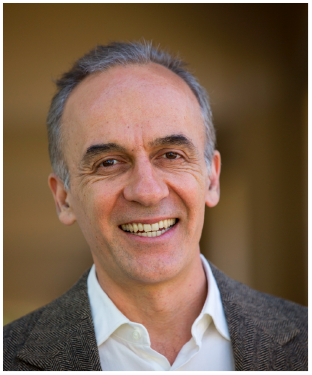

**E. J. Chichilnisky** is the John R. Adler Professor of Neurosurgery, and Professor of Ophthalmology, at Stanford University, where he has worked since 2013. Previously, he worked at the Salk Institute for Biological Studies for 15 years. He received his B.A. in Mathematics from Princeton University, and his M.S. in mathematics and Ph.D. in neuroscience from Stanford University. His research has focused on understanding the spatiotemporal patterns of electrical activity in the retina that convey visual information to the brain, and their origins in retinal circuitry, using large-scale multielectrode recordings. His ongoing work now focuses on using basic science knowledge along with electrical stimulation to develop a novel high-fidelity artificial retina for treating incurable blindness. He is the recipient of an Alfred P. Sloan Research Fellowship, a McKnight Scholar Award, a McKnight Technological Innovation in Neuroscience Award, and a Research to Prevent Blindness Stein Innovation Award.

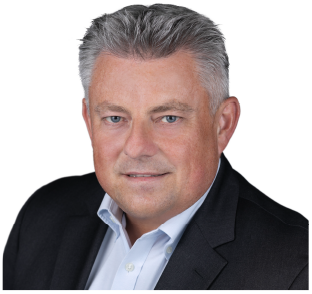

**Boris Murmann** (Fellow, IEEE) received the Dipl.-Ing. (FH) degree in communications engineering from Fachhochschule Dieburg, Dieburg, Germany, in 1994, the M.S. degree in electrical engineering from Santa Clara University, Santa Clara, CA, USA, in 1999, and the Ph.D. degree in electrical engineering from the University of California at Berkeley, Berkeley, CA, USA, in 2003.

From 1994 to 1997, he was with Neutron Mikroelektronik GmbH, Hanau, Germany, where he was involved in the development of low-power and smart-power application-specific integrated circuits (ASICs) in automotive CMOS technology. From 2004 to 2023, he was with the Department of Electrical Engineering, Stanford University, Stanford, CA, USA, where he served as an Assistant Professor, an Associate Professor, and a Full Professor. He is currently with the Department of Electrical and Computer Engineering, University of Hawai’i at Mānoa, Honolulu, HI, USA. His research interests are in the area of mixed-signal integrated circuit design, including sensor interfaces, A/D and D/A conversion, high-speed communication links, embedded machine learning (tinyML), electronic design automation, as well as open-source chip design.

Dr. Murmann was a co-recipient of the Best Student Paper Award at the Very Large-Scale Integration Circuits Symposium in 2008 and 2021 and the Best Invited Paper Award at the IEEE Custom Integrated Circuits Conference (CICC) in 2008. He received the Agilent Early Career Professor Award in 2009, the Friedrich Wilhelm Bessel Research Award in 2012, the SIA-SRC University Researcher Award for lifetime research contributions to the U.S. semiconductor industry in 2021, and the SRC Aristotle Award for contributions to teaching and mentorship in 2024. He was the 2017 Program Chair of the IEEE International Solid-State Circuits Conference (ISSCC) and the 2023 General Co-Chair of the IEEE International Symposium on Circuits and Systems (ISCAS). He currently serves as the Editor-in-Chief of the IEEE Journal of Solid-State Circuits.

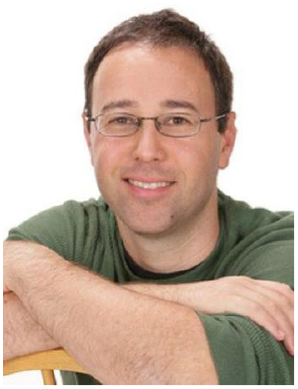

**Tsachy weissman** (Fellow, IEEE), is the Robert and Barbara Kleist Professor of Electrical Engineering at Stanford University since 2003, researching and teaching the science of information, with applications spanning genomics, neuroscience, and technology. He has been serving on editorial boards for scientific journals, technical advisory boards in industry, and as the Founding Director of the Stanford Compression Forum. His recent projects include the SHTEM summer internship program for high schoolers, the starling initiative for data integrity, and stagecast, a low-latency video platform allowing actors and singers to perform together in real-time while geographically distributed. He has received multiple awards for his research and teaching, including Best Paper Awards from the IEEE Information Theory and Communications Societies, while his students received best student authored paper awards at the top conferences of their areas of scholarship.

He has prototyped guardant health’s first algorithms for early detection of cancer from blood tests and has recently co-founded and sold compressable to Amazon, reducing humanity’s cloud storage footprint. His favorite gig to date was advising the HBO show “Silicon Valley.”

## Footnotes

1 Intuitively, the observed voltage at a given electrode is the superposition of the underlying spike signal and Gaussian noise. In such settings, quantization optimized for the clean signal often performs nearly as well as one optimized for the noisy observation. See [47] for theoretical discussion.

2 Here the mean square error is normalized to the spike peak to peak amplitude *Vpeak−peak amplitude*.

3 This is an empirical number dependent on the sampling rate of the system.

4 Electrical images are generated by averaging spike waveforms to capture the characteristic signature of each cell–electrode pair.

5 For each potential spike, we consider the center electrode and its 6 adjacent electrodes and buffer 30 samples per electrode.

6 A conservative 4KB estimate accounts for two 1024-entry Huffman lookup tables of the channel addresses and recorded quantized ADC amplitudes.

7 Here, we use the compression ratio results from the ExVivo-2 simulation as a representative example.

## References

[1] Y. Wang, X. Yang, X. Zhang, Y. Wang, and W. Pei, “Implantable intracortical microelectrodes: reviewing the present with a focus on the future,” Microsyst Nanoeng, vol. 9, p. 7, Jan. 2023.

[2] T. Matsuo, K. Kawai, T. Uno, N. Kunii, N. Miyakawa, K. Usami, K. Kawasaki, I. Hasegawa, and N. Saito, “Simultaneous recording of single-neuron activities and broad-area intracranial electroencephalography: electrode design and implantation procedure,” Neurosurgery, vol. 73, no. 2 Suppl Operative, pp. ons146–54, Dec. 2013.

[3] A. L. Juavinett, G. Bekheet, and A. K. Churchland, “Chronically implanted neuropixels probes enable high-yield recordings in freely moving mice,” eLife, vol. 8, Aug. 2019. [Online]. Available: https://doi.org/10.7554/elife.47188

[4] J. Putzeys, B. C. Raducanu, A. Carton, J. De Ceulaer, B. Karsh, J. H. Siegle, N. Van Helleputte, T. D. Harris, B. Dutta, S. Musa, and C. Mora Lopez, “Neuropixels Data-Acquisition system: A scalable platform for parallel recording of 10 000+ electrophysiological signals,” IEEE Trans. Biomed. Circuits Syst., vol. 13, no. 6, pp. 1635–1644, Dec. 2019.

[5] T. Z. Luo, A. G. Bondy, D. Gupta, V. A. Elliott, C. D. Kopec, and C. D. Brody, “An approach for long-term, multi-probe neuropixels recordings in unrestrained rats,” Elife, vol. 9, Oct. 2020.

[6] A. C. Paulk, Y. Kfir, A. R. Khanna, M. L. Mustroph, E. M. Trautmann, D. J. Soper, S. D. Stavisky, M. Welkenhuysen, B. Dutta, K. V. Shenoy, L. R. Hochberg, R. M. Richardson, Z. M. Williams, and S. S. Cash, “Large-scale neural recordings with single neuron resolution using neuropixels probes in human cortex,” Nat. Neurosci., vol. 25, no. 2, pp. 252–263, Feb. 2022.

[7] E. M. Trautmann, J. K. Hesse, G. M. Stine, R. Xia, S. Zhu, D. J. O’Shea, B. Karsh, J. Colonell, F. F. Lanfranchi, S. Vyas, A. Zimnik, N. A. Steinmann, D. A. Wagenaar, A. Andrei, C. M. Lopez, J. O’Callaghan, J. Putzeys, B. C. Raducanu, M. Welkenhuysen, M. Churchland, T. Moore, M. Shadlen, K. Shenoy, D. Tsao, B. Dutta, and T. Harris, “Large-scale high-density brain-wide neural recording in nonhuman primates,” bioRxiv, 2023, available at: https://www.biorxiv.org/content/early/2023/05/04/2023.02.01.526664. x[Online]. Available: https://www.biorxiv.org/content/early/2023/05/04/2023.02.01.526664

[8] F. R. Willett, E. M. Kunz, C. Fan, D. T. Avansino, G. H. Wilson, E. Y. Choi, F. Kamdar, M. F. Glasser, L. R. Hochberg, S. Druckmann, K. V. Shenoy, and J. M. Henderson, “A high-performance speech neuroprosthesis,” Nature, vol. 620, no. 7976, pp. 1031–1036, Aug 2023. [Online]. Available: https://doi.org/10.1038/s41586-023-06377-x

[9] P. D. Ganzer, S. C. Colachis, 4th, M. A. Schwemmer, D. A. Friedenberg, C. F. Dunlap, C. E. Swiftney, A. F. Jacobowitz, D. J. Weber, M. A. Bockbrader, and G. Sharma, “Restoring the sense of touch using a sensorimotor demultiplexing neural interface: ‘disentangling’ sensorimotor events during brain-computer interface control,” in SpringerBriefs in Electrical and Computer Engineering. Cham: Springer International Publishing, 2021, pp. 75–85.

[10] N. A. Steinmetz, C. Aydin, A. Lebedeva, M. Okun, M. Pachitariu, M. Bauza, M. Beau, J. Bhagat, C. Böhm, M. Broux, S. Chen, J. Colonell, R. J. Gardner, B. Karsh, F. Kloosterman, D. Kostadinov, C. Mora-Lopez, J. O’Callaghan, J. Park, J. Putzeys, B. Sauerbrei, R. J. J. van Daal, A. Z. Vollan, S. Wang, M. Welkenhuysen, Z. Ye, J. T. Dudman, B. Dutta, A. W. Hantman, K. D. Harris, A. K. Lee, E. I. Moser, J. O’Keefe, A. Renart, K. Svoboda, M. Häusser, S. Haesler, M. Carandini, and T. D. Harris, “Neuropixels 2.0: A miniaturized high-density probe for stable, longterm brain recordings,” Science, vol. 372, no. 6539, p. eabf4588, Apr. 2021.

[11] K. Sahasrabuddhe, A. A. Khan, A. P. Singh, T. M. Stern, Y. Ng, A. Tadić, P. Orel, C. LaReau, D. Pouzzner, K. Nishimura, K. M. Boergens, S. Shivakumar, M. S. Hopper, B. Kerr, M.-E. S. Hanna, R. J. Edgington, I. McNamara, D. Fell, P. Gao, A. Babaie-Fishani, S. Veijalainen, A. V. Klekachev, A. M. Stuckey, B. Luyssaert, T. D. Y. Kozai, C. Xie, V. Gilja, B. Dierickx, Y. Kong, M. Straka, H. S. Sohal, and M. R. Angle, “The argo: a high channel count recording system for neural recording in vivo,” J. Neural Eng., vol. 18, no. 1, p. 015002, Feb. 2021.

[12] E. Musk and Neuralink, “An integrated Brain-Machine interface platform with thousands of channels,” J. Med. Internet Res., vol. 21, no. 10, p. e16194, Oct. 2019.

[13] H. G. Rey, C. Pedreira, and R. Quian Quiroga, “Past, present and future of spike sorting techniques,” Brain Res Bull, vol. 119, no. Pt B, pp. 106–117, Apr. 2015.

[14] G. T. Einevoll, C. Kayser, N. K. Logothetis, and S. Panzeri, “Modelling and analysis of local field potentials for studying the function of cortical circuits,” Nat Rev Neurosci, vol. 14, no. 11, pp. 770–785, Nov. 2013.

[15] A. Nandi, T. Chartrand, W. Van Geit, A. Buchin, Z. Yao, S. Y. Lee, Y. Wei, B. Kalmbach, B. Lee, E. Lein, J. Berg, U. Sümbül, C. Koch, B. Tasic, and C. A. Anastassiou, “Single-neuron models linking electrophysiology, morphology, and transcriptomics across cortical cell types,” Cell Rep, vol. 40, no. 6, p. 111176, Aug. 2022.

[16] N. W. Gouwens, S. A. Sorensen, J. Berg, C. Lee, T. Jarsky, J. Ting, S. M. Sunkin, D. Feng, C. A. Anastassiou, E. Barkan, K. Bickley, N. Blesie, T. Braun, K. Brouner, A. Budzillo, S. Caldejon, T. Casper, D. Castelli, P. Chong, K. Crichton, C. Cuhaciyan, T. L. Daigle, R. Dalley, N. Dee, T. Desta, S.-L. Ding, S. Dingman, A. Doperalski, N. Dotson, T. Egdorf, M. Fisher, R. A. de Frates, E. Garren, M. Garwood, A. Gary, N. Gaudreault, K. Godfrey, M. Gorham, H. Gu, C. Habel, K. Hadley, J. Harrington, J. A. Harris, A. Henry, D. Hill, S. Josephsen, S. Kebede, L. Kim, M. Kroll, B. Lee, T. Lemon, K. E. Link, X. Liu, B. Long, R. Mann, M. McGraw, S. Mihalas, A. Mukora, G. J. Murphy, L. Ng, K. Ngo, T. N. Nguyen, P. R. Nicovich, A. Oldre, D. Park, S. Parry, J. Perkins, L. Potekhina, D. Reid, M. Robertson, D. Sandman, M. Schroedter, C. Slaughterbeck, G. Soler-Llavina, J. Sulc, A. Szafer, B. Tasic, N. Taskin, C. Teeter, N. Thatra, H. Tung, W. Wakeman, G. Williams, R. Young, Z. Zhou, C. Farrell, H. Peng, M. J. Hawrylycz, E. Lein, L. Ng, A. Arkhipov, A. Bernard, J. W. Phillips, H. Zeng, and C. Koch, “Classification of electrophysiological and morphological neuron types in the mouse visual cortex,” Nat Neurosci, vol. 22, no. 7, pp. 1182–1195, Jun. 2019.

[17] O. Ophir, O. Shefi, and O. Lindenbaum, “Classifying neuronal cell types based on shared electrophysiological information from humans and mice,” Neuroinformatics, vol. 22, no. 4, pp. 473–486, Oct 2024. [Online]. Available: 10.1007/s12021-024-09675-5

[18] E. M. Trautmann, S. D. Stavisky, S. Lahiri, K. C. Ames, M. T. Kaufman, D. J. O’Shea, S. Vyas, X. Sun, S. I. Ryu, S. Ganguli, and K. V. Shenoy, “Accurate estimation of neural population dynamics without spike sorting,” Neuron, vol. 103, no. 2, pp. 292–308.e4, Jul. 2019.

[19] N. Even-Chen, D. G. Muratore, S. D. Stavisky, L. R. Hochberg, J. M. Henderson, B. Murmann, and K. V. Shenoy, “Power-saving design opportunities for wireless intracortical brain–computer interfaces,” Nature Biomedical Engineering, vol. 4, no. 10, pp. 984–996, Aug. 2020.

[20] S.-Y. Park, J. Cho, K. Lee, and E. Yoon, “Dynamic Power Reduction in Scalable Neural Recording Interface Using Spatiotemporal Correlation and Temporal Sparsity of Neural Signals,” IEEE journal of solid-state circuits, vol. 53, no. 4, pp. 1102–1114, Apr. 2018. [Online]. Available: http://dx.doi.org/10.1109/JSSC.2017.2787749

[21] X. Guo, M. Shaeri, and M. Shoaran, “An Accurate and Hardware-Efficient Dual Spike Detector for Implantable Neural Interfaces,” in 2022 IEEE Biomedical Circuits and Systems Conference (BioCAS), Oct. 2022, pp. 70–74. [Online]. Available: http://dx.doi.org/10.1109/BioCAS54905.2022.9948602

[22] M. A. Shaeri and A. M. Sodagar, “A Method for Compression of Intra-Cortically-Recorded Neural Signals Dedicated to Implantable Brain–Machine Interfaces,” IEEE transactions on neural systems and rehabilitation engineering: a publication of the IEEE Engineering in Medicine and Biology Society, vol. 23, no. 3, pp. 485–497, May 2015. [Online]. Available: http://dx.doi.org/10.1109/TNSRE.2014.2355139

[23] V. Mohan, W. P. Tay, and A. Basu, “Towards neuromorphic compression based neural sensing for next-generation wireless implantable brain machine interface,” Neuromorphic Computing and Engineering, vol. 5, no. 1, p. 014004, jan 2025. [Online]. Available: https://dx.doi.org/10.1088/2634-4386/adad10

[24] D. Valencia and A. Alimohammad, “A Real-Time spike sorting system using parallel OSort clustering,” IEEE Trans. Biomed. Circuits Syst., vol. 13, no. 6, pp. 1700–1713, Dec. 2019.

[25] Y. Chen, B. Tacca, Y. Chen, D. Biswas, G. Gielen, F. Catthoor, M. Verhelst, and C. M. Lopez, “A 384-Channel Online-Spike-Sorting IC Using Unsupervised Geo-OSort Clustering and Achieving 0.0013mm2/Ch and 1.78µW/ch,” in 2023 IEEE International Solid-State Circuits Conference (ISSCC), Feb. 2023, pp. 486–488. [Online]. Available: http://dx.doi.org/10.1109/ISSCC42615.2023.10067264

[26] J. Li, P. K. Chundi, S. Kim, Z. Jiang, M. Yang, J. Kang, S. Jung, S. J. Kim, and M. Seok, “A 0.78-μW 96-Ch. Deep Sub-Vt Neural Spike Processor Integrated with a Nanowatt Power Management Unit,” in ESSCIRC 2018 -IEEE 44th European Solid State Circuits Conference (ESSCIRC), Sep. 2018, pp. 154–157. [Online]. Available: http://dx.doi.org/10.1109/ESSCIRC.2018.8494273

[27] T. Wu, W. Zhao, E. Keefer, and Z. Yang, “Deep compressive autoencoder for action potential compression in large-scale neural recording,” Journal of Neural Engineering, vol. 15, no. 6, p. 066019, oct 2018. [Online]. Available: https://doi.org/10.1088%2F1741-2552%2Faae18d

[28] M. Pagin and M. Ortmanns, “A neural data lossless compression scheme based on spatial and temporal prediction,” in 2017 IEEE Biomedical Circuits and Systems Conference (BioCAS). IEEE, Oct. 2017.

[29] J. Thies and A. Alimohammad, “Compact and low-power neural spike compression using undercomplete autoencoders,” IEEE Transactions on Neural Systems and Rehabilitation Engineering, vol. 27, no. 8, pp. 1529– 1538, 2019.

[30] A. Kipnis, Y. C. Eldar, and A. J. Goldsmith, “Analog-to-Digital compression: A new paradigm for converting signals to bits,” IEEE Signal Process. Mag., vol. 35, no. 3, pp. 16–39, May 2018.

[31] A. Akhoundi, Y. Landbrug, P. Yan, E. J. Chichilnisky, B. Murmann, and D. G. Muratore, “15.2 a 1024-channel 0.00029mm2/ch 74nw/ch online spatial spike-sorting chip with event-driven spike detection and self-organizing map clustering,” in 2025 IEEE International Solid-State Circuits Conference (ISSCC), vol. 68, 2025, pp. 268–270.

[32] Y. Chen, B. Tacca, Y. Chen, D. Biswas, G. Gielen, F. Catthoor, M. Verhelst, and C. Mora Lopez, “An online-spike-sorting ic using unsupervised geometry-aware osort clustering for efficient embedded neural-signal processing,” IEEE Journal of Solid-State Circuits, vol. 58, no. 11, pp. 2990–3002, 2023.

[33] D. G. Muratore, P. Tandon, M. Wootters, E. J. Chichilnisky, S. Mitra, and B. Murmann, “A data-compressive wired-or readout for massively parallel neural recording,” IEEE Transactions on Biomedical Circuits and Systems, vol. 13, no. 6, pp. 1128–1140, 2019.

[34] P. Yan, A. Akhoundi, N. P. Shah, P. Tandon, D. G. Muratore, E. J. Chichilnisky, and B. Murmann, “Data compression versus signal fidelity tradeoff in wired-or analog-to-digital compressive arrays for neural recording,” IEEE Transactions on Biomedical Circuits and Systems, vol. 17, no. 4, pp. 754–767, 2023.

[35] M. Jang, M. Hays, W.-H. Yu, C. Lee, P. Caragiulo, A. T. Ramkaj, P. Wang, A. J. Phillips, N. Vitale, P. Tandon, P. Yan, P.-I. Mak, Y. Chae, E. J. Chichilnisky, B. Murmann, and D. G. Muratore, “A 1024-channel 268-nw/pixel 36×36 um2/channel data-compressive neural recording ic for high-bandwidth brain–computer interfaces,” IEEE Journal of SolidState Circuits, vol. 59, no. 4, pp. 1123–1136, 2024.

[36] J. J. Jun, N. A. Steinmetz, J. H. Siegle, D. J. Denman, M. Bauza, B. Barbarits, A. K. Lee, C. A. Anastassiou, A. Andrei, Ç. Aydin, M. Barbic, T. J. Blanche, V. Bonin, J. Couto, B. Dutta, S. L. Gratiy, D. A. Gutnisky, M. Häusser, B. Karsh, P. Ledochowitsch, C. M. Lopez, C. Mitelut, S. Musa, M. Okun, M. Pachitariu, J. Putzeys, P. D. Rich, C. Rossant, W.-L. Sun, K. Svoboda, M. Carandini, K. D. Harris, C. Koch, J. O’Keefe, and T. D. Harris, “Fully integrated silicon probes for high-density recording of neural activity,” Nature, vol. 551, no. 7679, pp. 232–236, Nov. 2017.

[37] J. L. Shobe, L. D. Claar, S. Parhami, K. I. Bakhurin, and S. C. Masmanidis, “Brain activity mapping at multiple scales with silicon microprobes containing 1,024 electrodes,” J. Neurophysiol., vol. 114, no. 3, pp. 2043–2052, Sep. 2015.

[38] B. J. Black, A. Kanneganti, A. Joshi-Imre, R. Rihani, B. Chakraborty, J. Abbott, J. J. Pancrazio, and S. F. Cogan, “Chronic recording and electrochemical performance of utah microelectrode arrays implanted in rat motor cortex,” J. Neurophysiol., vol. 120, no. 4, pp. 2083–2090, Oct. 2018.

[39] R. Bartolo, R. C. Saunders, A. R. Mitz, and B. B. Averbeck, “Dimensionality, information and learning in prefrontal cortex,” PLoS Comput. Biol., vol. 16, no. 4, p. e1007514, Apr. 2020.

[40] M. Pachitariu, S. Sridhar, J. Pennington, and C. Stringer, “Spike sorting with kilosort4,” Nature Methods, vol. 21, no. 5, pp. 914–921, May 2024. [Online]. Available: https://doi.org/10.1038/s41592-024-02232-7

[41] J. Ziv, “On universal quantization,” IEEE Transactions on Information Theory, vol. 31, no. 3, pp. 344–347, 1985.

[42] H. Witsenhausen, “Indirect rate distortion problems,” IEEE Transactions on Information Theory, vol. 26, no. 5, pp. 518–521, 1980.

[43] T. Weissman and E. Ordentlich, “The empirical distribution of rate-constrained source codes,” in International Symposium onInformation Theory, 2004. ISIT 2004. Proceedings., 2004, pp. 464–.

[44] S. Jalali and T. Weissman, “Denoising via MCMC-based lossy compression,” IEEE Transactions on Signal Processing, vol. 60, no. 6, pp. 3092–3100, 2012.

[45] I. Ochoa, M. Hernaez, R. Goldfeder, T. Weissman, and E. Ashley, “Effect of lossy compression of quality scores on variant calling,” Brief Bioinform, vol. 18, no. 2, pp. 183–194, Mar. 2017.

[46] B. Natarajan, K. Konstantinides, and C. Herley, “Occam filters for stochastic sources with application to digital images,” IEEE Transactions on Signal Processing, vol. 46, no. 5, pp. 1434–1438, 1998.

[47] Y. Ephraim and R. Gray, “A unified approach for encoding clean and noisy sources by means of waveform and autoregressive model vector quantization,” IEEE Transactions on Information Theory, vol. 34, no. 4, pp. 826–834, 1988.

[48] “Neuropixels datasets, ‘sorting comparison results’,” http://phy.cortexlab.net/data/sortingComparison/, 2016, xaccessed: May 10, 2023.

[49] S. E. Paraskevopoulou, D. Wu, A. Eftekhar, and T. G. Constandinou, “Hierarchical adaptive means (ham) clustering for hardware-efficient, unsupervised and real-time spike sorting,” Journal of Neuroscience Methods, vol. 235, pp. 145–156, 2014. [Online]. Available: https://www.sciencedirect.com/science/article/pii/S0165027014002489

[50] P. Yan, N. P. Shah, D. G. Muratore, P. Tandon, E. Chichilnisky, and B. Murmann, “Data compression versus signal fidelity trade-off in wiredor adc arrays for neural recording,” in 2022 IEEE Biomedical Circuits and Systems Conference (BioCAS), 2022, pp. 80–84.

[51] A. Pedram, S. Richardson, M. Horowitz, S. Galal, and S. Kvatinsky, “Dark memory and accelerator-rich system optimization in the dark silicon era,” IEEE Design & Test, vol. 34, no. 2, pp. 39–50, 2017.

[52] K. Prabhu, R. M. Radway, J. Yu, K. Bartolone, M. Giordano, F. Peddinghaus, Y. Urman, W.-S. Khwa, Y.-D. Chih, M.-F. Chang, S. Mitra, and P. Raina, “Minotaur: A posit-based 0.42–0.50-tops/w edge transformer inference and training accelerator,” IEEE Journal of Solid-State Circuits, vol. 60, no. 4, pp. 1311–1323, 2025.

[53] D. Harris and S. Harris, Digital Design and Computer Architecture, Second Edition, 2nd ed. San Francisco, CA, USA: Morgan Kaufmann Publishers Inc., 2012.

[54] V. K. Kodavalla, “Ip gate count estimation methodology during micro-architecture phase,” Design & Reuse, September 2008, accessed: 2025-04-23. [Online]. Available: https://www.design-reuse.com/articles/19171/ip-gate-count-estimation-micro-architecture-phase.html

[55] E. So and A. Arbabian, “6.1 12mb/s 4×4 ultrasound mimo relay with wireless power and communication for neural interfaces,” in 2024 IEEE International Solid-State Circuits Conference (ISSCC), vol. 67, 2024, pp. 100–102.

[56] M. Aboelmaged, A. Shisha, and M. A. A. E. Ghany, “High-performance data compression-based design for dynamic iot security systems,” Electronics, vol. 10, no. 16, 2021. [Online]. Available: https://www.mdpi.com/2079-9292/10/16/1989

